# Resilience to anxiety and anhedonia after predator scent stress is accompanied by increased nucleus accumbens mGlu5 in female rats

**DOI:** 10.1101/2022.04.15.488499

**Authors:** H.L. Blount, J. Dee, L. Wu, M. Schwendt, L.A. Knackstedt

## Abstract

Despite the higher prevalence of post-traumatic stress disorder (PTSD) in women, the majority of preclinical neuroscience research has been conducted utilizing male subjects. We have found that male rats exposed to the predator scent 2,4,5-trimethyl-3-thiazoline (TMT) show heterogenous development of long-term anxiety-like behavior and conditioned fear to the TMT environment. Stress-Resilient males exhibit increased mGlu5 mRNA expression in the basolateral amygdala (BLA) and prefrontal cortex. Here we sought to determine whether the same behavioral and genetic responses would be observed in female rats exposed to TMT. Sprague-Dawley rats were exposed to TMT for ten minutes, while Controls were exposed to an unscented environment. Anxiety and anhedonia were assessed 7-14 days later with elevated plus maze (EPM), acoustic startle response (ASR), light/dark box, and sucrose preference test. TMT-exposed females spent less time in the EPM open arms and exhibited greater startle amplitude, and reduced sucrose intake compared to Controls. Median split analyses conducted on EPM and sucrose intake yielded phenotypes that displayed behavior in the light/dark box consistent with EPM and sucrose testing. Unlike male Susceptible rats, female Susceptible rats showed no freezing when re-exposed to the TMT context, nor did Resilient female rats present elevated BLA mGlu5 mRNA levels. Instead, Susceptible females had greater BLA mGlu5 than Resilient or Control rats. This work indicates that, as in humans, rats exhibit sex-dependent responses to stress. This translational animal model may provide insight into how females are uniquely affected by PTSD.

## Introduction

Post-traumatic stress disorder (PTSD) is an anxiety disorder that develops only in a subpopulation (15-25%) of trauma-affected individuals (1–3). Epidemiological studies report lifetime PTSD rates of approximately 6.2%-8.2% for men and 13.0%-20.4% for women (3–5). While there are behavioral and pharmacotherapeutic treatments for PTSD, these are not effective in all patients. Further exploration of the biological systems mediating anhedonia, anxiety, fear, and fear memory is necessary to improve existing therapeutics and identify novel treatment options for trauma-associated psychopathology.

Animal models are fundamental for elucidating the neurobiology of PTSD susceptibility and resilience (6–9). One approach to developing preclinical models of PTSD is to back-translate criterion A of the DSM-5’s diagnostic criteria: an acute sense of endangerment and helplessness. Animal models of PTSD with face validity include chronic foot shock, forced swim, submersion, or physical immobilization (10,11). Predator or predator scent exposure are models of inescapable stress that offer additional ethological validity. In these paradigms, rodents are placed in an enclosure with a nearby live feline or the scent of a predator such as feline urine or the synthetic fox pheromone 2,5-dihydro-2,4,5-trimethylthiazoline (TMT) (9,12,13). Our lab has previously demonstrated that outbred male rats confined to a chamber with TMT predator scent stress (PSS) for 10 minutes demonstrate heterogeneous severity of anxiety-like behavior 7 days after exposure and context-induced fear (freezing) 21 days after exposure. In agreement with others who have used a similar model employing cat urine as the stressor, our group has found that approximately 25% of male rats display behavior in the elevated plus maze (EPM) and acoustic startle response (ASR) that is no different from that exhibited by non-TMT-exposed controls, thus demonstrating “resilience” to the stressor (14,15). Another 20% - 25% of “Susceptible” rats fall on the opposite end of the spectrum, showing anxiety and contextual fear that is significantly greater than observed from either controls or Resilient rats. In this project, we aim to adapt our model to research PTSD using female subjects.

Clinical and animal research identified that reciprocal mPFC-BLA circuits play a role in anxiety, fear memory -formation, -maintenance, and -extinction (16–20). While several neurotransmitter systems have been implicated in pathological fear and anxiety, dysregulation of glutamatergic tone within this circuit has been documented in PTSD and other anxiety disorders (21– 25). Critically, glutamate levels in the prefrontal cortex predict the magnitude of anxiety responses in both healthy and trauma-exposed subjects (26,27). Metabotropic glutamate receptor 5 (mGlu5) has been implicated in mediating resilience to stress-induced anhedonia and anxiety in rodents (28–32). In agreement with those works, we have found that male rats “resilient” to the long-term effects of predator scent stress (PSS) exhibit greater PFC and BLA mGlu5 mRNA expression, and mGlu5 antagonists and agonists bidirectionally alter contextual fear in this model (31,33–37).

Despite the prevalence and increased risk for the development of PTSD in women, few preclinical studies have examined stress-susceptibility or its neurobiological underpinnings utilizing female rodents (38–40). The current study seeks to determine if outbred female rats exhibit anxiety-like behavior and fear following a single TMT exposure in a similar manner as male rats. Trauma susceptibility may manifest itself more differently in female rats than observed in male rats. Because there is evidence that women with PTSD show more symptoms of anhedonia than men, we hypothesized that stress-Susceptible female rats would show anhedonia (36). We also examined whether Resilient female rats would express increased PFC and BLA mGlu5 mRNA, akin to male rats. Because mGlu5 expression in the nucleus accumbens (NAc) has been linked with resilience against the development of stress-induced anhedonia, we hypothesized that Resilient female rats would exhibit greater mGlu5 in this brain region (28,29). We also hypothesized that there may be a possible role for the estrous cycle at the time of predator scent stress or anxiety testing in influencing individual differences of vulnerability to developing PTSD-like behaviors.

## 2. Materials & Methods

### 2.1 Subjects

Sprague-Dawley female rats (Charles River Laboratories, Raleigh, NC, USA) were individually housed in climate-regulated cages on a 12-h reverse light cycle (7am lights off). Rats were 8 weeks of age at arrival and acclimated to the vivarium for 7 days with ad libitum access to water and chow. Five days prior to TMT or control exposure, subjects were food restricted to 20 g/day and acclimated to handling and vaginal lavage. All animal procedures for behavioral and biochemical experiments were authorized by University of Florida’s Institutional Animal Care & Use Committee (IACUC). One female and one male experimenter conducted all procedures within four hours of the beginning of dark cycle. Morning experimental timing may have influenced the distribution of estrus-staging at TMT-exposure and EPM-ASR testing, skewing our findings towards di-, pro-, and meta-estrus staging. Fifty-three rats were used for these experiments.

### 2.2 Predator scent stress (PSS)

Predator scent and control exposures occurred in separate circular plexiglass chambers 35cm high and 40 cm in diameter (Bio Bubble Pets, Boca Raton, FL, USA). Steel mesh flooring ensured olfactory exposure without direct contact with the predator pheromone. Two subjects were exposed at a time in adjacent, individual, clear plexiglass containers. Rats were either exposed to an unscented chamber (n=13) or one containing TMT (n=40). During TMT exposure, 3 uL of TMT (>97% purity; BioSRQ, Sarasota, FL, USA) was blotted onto paper and placed underneath the stainless-steel mesh floor (34). Control rats were exposed to a no-scent condition in a separate set of plexiglass chambers that never contained TMT. Exposures were recorded by an external HD camera to measure freezing, rearing, center crossing, and grooming behavior. Prior to and between each exposure session, equipment was cleaned using 70% EtOH and aerated to eliminate lingering EtOH odor.

### 2.3 Elevated Plus Maze

One week following TMT/control exposure (see Fig. 1A), rats were evaluated for anxiety-like behavior using an elevated plus maze (EPM). The black acrylic maze was suspended 50 cm off the ground, and arms were 50 cm x 10 cm joined by a 10 cm square center platform. The walls of the two closed arms were 50 cm high, while the two open arms’ walls were 2.5 cm in height. The EPM was lit with diffused light (∼ 50 lux). Rats were placed into the center platform and the test lasted 5 min. The maze was stationed under a camera for HD video recording. EthoVision XT 15 software (Noldus Information Technology) was used to quantify time spent in the open arms (OA) and closed arms (CA) and center platform, and number of entries into each arm. Prior to each testing session, the maze was cleaned using 70% EtOH.

**Figure 1.**
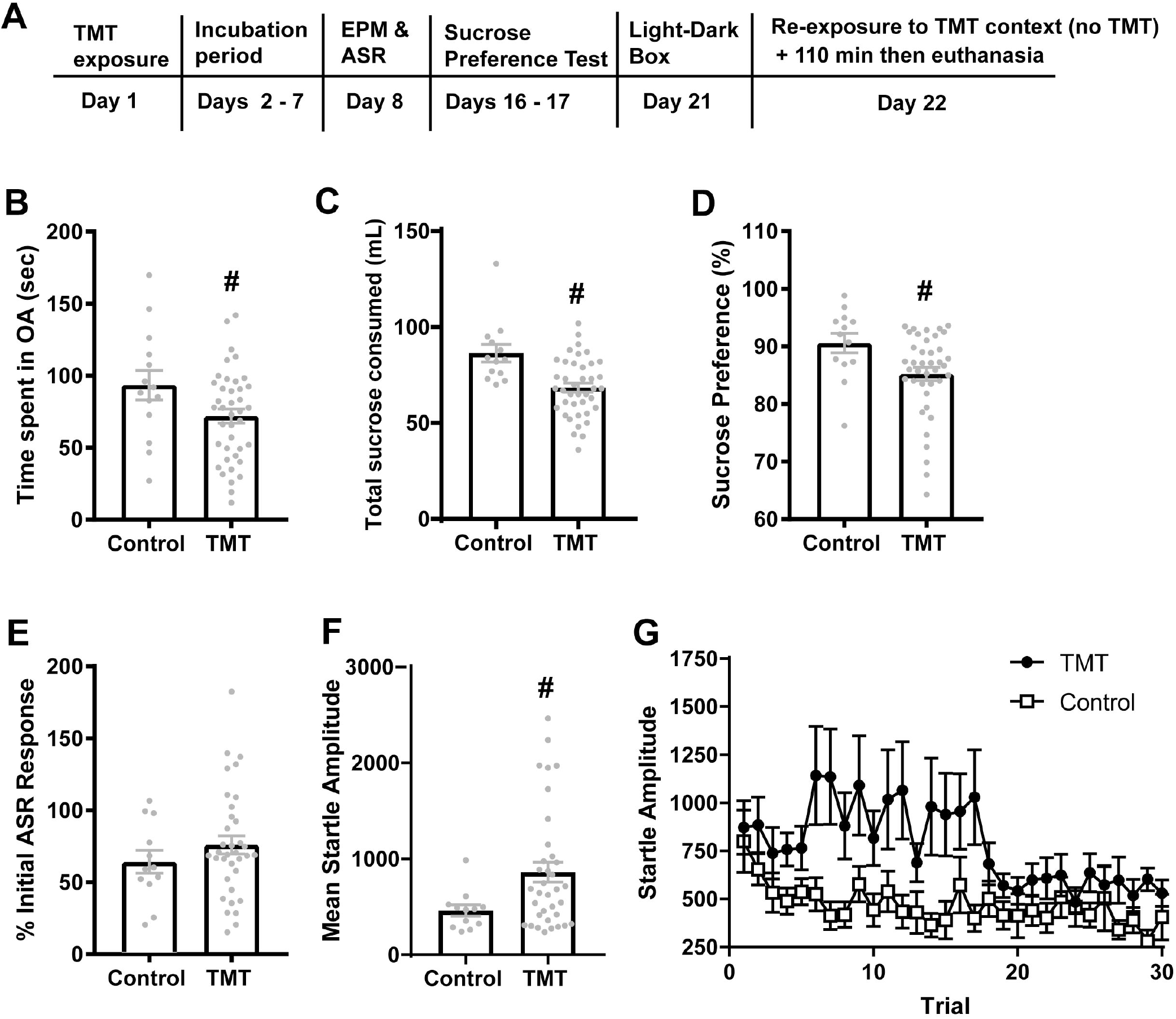
Experimental timeline and impact of TMT exposure on anxiety- and anhedonia-like behaviors. **(A)** Timeline of experiment. **(B)** Main effects of group [Control vs. TMT-exposed;] were observed for time spent in the Open Arms (OA) of the elevated plus maze (EPM), **(C)** total sucrose consumption, and **(D)** sucrose preference **(E)** Habituation of the acoustic startle response (ASR) did not differ between groups. **(F)** TMT-exposed subjects exhibited greater mean amplitude of the ASR than unstressed Controls. **(G)** Over 30 trials, ASR did not differ significantly between groups. TMT-exposed females exhibited elevated ASR in early session but habituated their ASR to that of controls within the 30-trial testing session. # = p<0.05 vs. Control.

### 2.4. Acoustic Startle

Within 20 minutes of completing the EPM test, rats were tested for acoustic startle response (ASR) inside ventilated, soundproof, chambers containing a transparent plexiglass cylinder resting atop a pressure-sensitive platform (San Diego Instruments, San Diego, CA). The ASR test began with a 5m acclimation period followed by 30 trials during which 110dB white noise was presented for 40ms followed by a randomized 30s-45s intertrial interval. Between each testing session equipment was cleaned using 70% EtOH.

### 2.5 Sucrose preference test

Sixteen days following TMT/control exposure, anhedonia-like behavior was measured in the home cage with a sucrose preference test (Fig. 1A). Animals were habituated to a unique 50 mL bottle containing water for 24 hours. The following days, two 50 mL bottles of the same design were presented in home cages, one containing water and the other 32% sucrose-water solution. Every 12 hours liquid consumption was recorded and bottles were refilled. The location of the bottles was reversed after the first 24 hours.

### 2.6 Light-dark box test

Twenty-one days following TMT/control exposure, rats underwent a light-dark box test to assess phenotypic differences in unconditioned fear, based on previous published methods (34,41). Our light-dark box apparatus consisted of two connected chambers of equal size (40 cm × 44 cm × 37 cm). The light-chamber had clear walls, no roofing, and was illuminated at approximately 300 lux. The dark chamber had opaque walls and was covered. The two were joined by a 5 cm rectangular cut out in the light-chamber facing wall of the dark chamber. In the beginning of light-dark box testing, animals were placed in the center of the box and allowed to roam uninterrupted for 10 minutes. Testing sessions were recorded, and subjects were manually measured on: latency to enter the dark box, latency to return to the light chamber upon first entering of the dark chamber, and overall time spent in the dark and light chambers.

### 2.7 Contextual fear

One week following sucrose preference test animals were reintroduced to the TMT context for a contextual fear test (see Fig. 1A). Control chambers that were identical to exposure chambers but never contained TMT were used for this test. Rats were placed into the apparatus for 5 min and behavior was recorded using an external HD camera. Ethovision XT 15 software was used to track freezing, rearing, center crossing, and grooming behavior. All experimental areas and surfaces were deep cleaned with 30% EtOH between each exposure session.

### 2.8 Vaginal lavage

Estrous samples were collected by gently washing the vaginal canal with phosphate buffered solution and air-drying the wash product onto glass slides. Samples were stained with 0.1% crystal violet solution for one-minute, followed by ddH2O wash. Air-dried cells were visualized with 20x bright-field microscope to assess the estrous stage. Cornified squamous epithelial cells, leukocytes, and/or nucleated epithelial cell ratios were assessed to determine the estrous stage (42–45). Following two days of habituation to the technique, vaginal lavage samples were collected immediately following predator scent exposure and ASR testing. Due to loss of samples, estrous lavages, on the day of EPM-ASR, were only obtained from 8 of the 14 controls.

### 2.9 Fluorescent in situ hybridization (FISH) for mGlu5 quantification

#### 2.9.1 Tissue processing and FISH procedures

2h following contextual fear testing, rats were euthanized by rapid decapitation, brains were dissected along the longitudinal fissure into hemispheres, flash frozen in isopentane, cooled on dry ice, and preserved at -80 °C. The tissue mRNA analysis was conducted in 1 hemisphere from 7-8 animals per phenotype with brains selected at random (Control n=8; Resilient n=8; Susceptible n=7). Serial coronal sections (12μm thick) containing the prefrontal cortex (3.24 to 2.52 mm relative to Bregma), nucleus accumbens (2.52 mm to 1.80 mm relative to Bregma), and basolateral amygdala/nucleus reuniens (−1.92 to -2.16 mm relative to Bregma) were collected using a freezing cryostat (Leica CM1950). Tissue slices were mounted onto Superfrost Plus Gold glass slides, air-dried at room-temperature for 20 minutes, and then stored at -80°C.

FISH was performed using a RNAscope Multiplex Fluorescent Reagent Kit according to manufacturer’s instructions [Advanced Cell Diagnostics (ACD); Newark, CA, USA]. Slide-mounted frozen tissue was fixed for 15 minutes in 4% paraformaldehyde (PFA; 4°C, pH 7.45), and then dehydrated in an ethanol gradient (50%, 70%, 100%). Slides were air-dried and then RT Protease IV (ACD, Cat. #322340) was added to sections to incubate in a HybEZ oven for 30 minutes (ACD, Cat. #321461). Post-protease digestion, section slides were twice washed with 1X PBS. Oligonucleotide single strand riboprobes were then prepared at a volume ratio of 1:50 (C1:C2). mRNA probe hybridization and signal amplification were performed for mGlu5 and vGluT1. The following Advanced Cell Diagnostics probes were utilized: GRM5 in channel 1 (ACD Cat. #471241) SLC17A7 in channel 2 (Cat. #317001). Probes were mixed at a volume ratio of 1:50 (C1:C2) applied to each coronal section and incubated in a HybEZ oven for 2 hours at 40°C. Following incubation, sections were placed into a vertical slide rack and washed twice in buffer solution (ACD Cat. #320058) for two minutes with gentle agitation. Cell nuclei were DAPI counterstained, and slides were cover slipped with ProLong Gold antifade mountant (ThermoFisher Scientific). Mounted slides were stored in the dark at 4°C to dry for 1-2 days. Clear nail polish was used to seal and preserve slides.

#### 2.5.2 Acquisition & quantification of FISH signal

Quantification of florescent optical density was done using 40X 16-bit grayscale images acquired using a sCMOS K5 camera mounted onto a Leica DM6b (software v. 2.41) epiflorescent microscope. All images were captured using a dry HC PL APO 40x/0.95 CORR objective (Leica Cat. #11506414). 10 um image stacks were taken at 1 um steps using Leica Application Suite X software (v. 3.7.423463). Images from the Z-step with greatest mRNA resolution were utilized for quantitative analyses. Two consecutively sliced tissue sections were imaged from each brain region.

CellProfiler software (v. 4.0.7) was utilized to quantify gene expression for mGlu5 and vGluT1. DAPI was used to label cell nuclei. Using DAPI-directed ROIs, mGlu5 and vGluT1 mRNA expression was quantified within an expanded nuclei region to measure synaptic and peri-synaptic ribonucleotide transcripts. Signal was artificially boosted using Speckles enhancement. Only mRNA masked within DAPI- and vGluT1-positive neurons were reported for statistical purposes (33).

### 2.6 Statistical analysis

PRISM (v.9.3.1, GraphPad, La Jolla, CA) software was used for all frequentist statistical analysis (alpha level set at p<0.05). First, unpaired t-tests were used to compare phenotypic differences in EPM variables, mean startle amplitude, and sucrose intake. Next, rats were categorized into groups as stress-Susceptible, Resilient, or Intermediate phenotypes using a double median split of time spent in the OA of the EPM and sucrose preference. One-way ANOVA and unpaired t-tests were used to compare phenotypic differences in EPM, ASR, sucrose preference, light-dark box testing, estrous staging, and subregion comparison of mRNA expression between treatment groups. Freezing, rearing, center crossing, grooming during TMT-exposure and contextual re-exposure were compared with two-way fixed factorial repeated measures ANOVA with Group as the between-subjects factor and Time as a within-subjects factor. Unpaired t-test significant interactions were investigated with multiple comparisons adjusted Tukey’s post hoc tests. One- and two-way ANOVA significant interactions were followed by Holm-Šídák’s multiple comparisons testing to determine effect directionality.

## 3. Results

### 3.1 Comparisons between TMT-exposed and Control rats

#### 3.1.1 Anxiety-like behavior

We found that TMT-exposed rats spent less time in the OA of the EPM relative to control rats [t(52)= 2.072, p=0.043; Fig. 1B]. There were no significant differences between TMT-exposed and control rats for time spent in the CA, number of CA entries, number of OA entries or time spent in the center. However, when adding together time spent in the center and time spent in CA, the TMT-exposed rats spent more time in these areas than the control rats [t(52)= 2.072, p=0.0433, not shown].

Due to ASR equipment malfunction, startle data from 6 TMT-exposed rats was corrupted and could not be used for analysis. When comparing the remaining 34 TMT-exposed rats to the 13 controls, there were no group differences in the habituation of the startle response [Fig. 1E]. However, there was a significant increase in mean startle response over all 30 trials in TMT-exposed rats compared to controls [t(45)=2.168, p =0.0355, Fig. 1D]. When examining startle responses over the entire 30 trials, there was no Group x Trial interaction and no main effect of Trial, but there was a trend for a main effect of Group [F(1, 51) = 3.64, p=0.062, Fig. 1G].

#### 3.1.2 Anhedonia

TMT-exposed rats consumed less sucrose (mL) than control rats [t(51)=3.624, p=0.007; Fig. 1C]. There was a significant reduction in the percent liquid consumed that was sucrose in the TMT-exposed rats [t(51)=2.404; Fig. 1D). The total amount of water consumed did not differ between groups (not shown).

### 3.2 Segregation into stress-susceptible and -resilient phenotypes

Because there were no effects of TMT exposure on habituation of the startle response, and because we were missing startle data from 6 rats, to segregate rats into stress-Susceptible and Resilient phenotypes, a double median split was conducted on sucrose consumed and time spent in the OA. The median sucrose intake was 67.25 mL and median time spent in the OA was 73.7 seconds. Using these criteria, TMT-exposed rats were phenotyped as stress-Susceptible (n=11) if they scored below the median time spent in the OA and below median sucrose intake; rats were phenotyped as stress-Resilient (n=12) if they scored above the median time spent in the OA and above median sucrose intake (Fig. 2A). The remaining rats (n=17) were categorized as Intermediate and eliminated from the experiment. The same criteria were applied to Control rats, with 11 of 13 Control rats meeting criteria for “Resilience”, and 2 rats showing “Intermediate” phenotype (Fig. 2B). We next compared time spent in the OA, sucrose intake, and mean ASR between Susceptible, Resilient and Control rats. There was a significant effect of Phenotype for time spent in the OA [F(2, 33) = 15.03, p<0.0001; Fig. 2C]. Post-hoc tests showed that Susceptible rats spent less time in the OA than both Control and Resilient rats (p’s <0.05). For total sucrose consumed (mL), there was a significant effect of Phenotype [F(2, 33) = 16.34, p<0.0001, Fig. 2D]. Post-hoc tests showed that Susceptible rats consumed less sucrose solution than both Control and Resilient rats (p’s <0.05). For mean ASR, there was a significant effect of Phenotype [F (2, 28) = 3.485, p=0.0445, Fig. 2E]. Post-hoc tests found that Susceptible rats displayed a mean startle response that was greater than Control rats.

**Figure 2.**
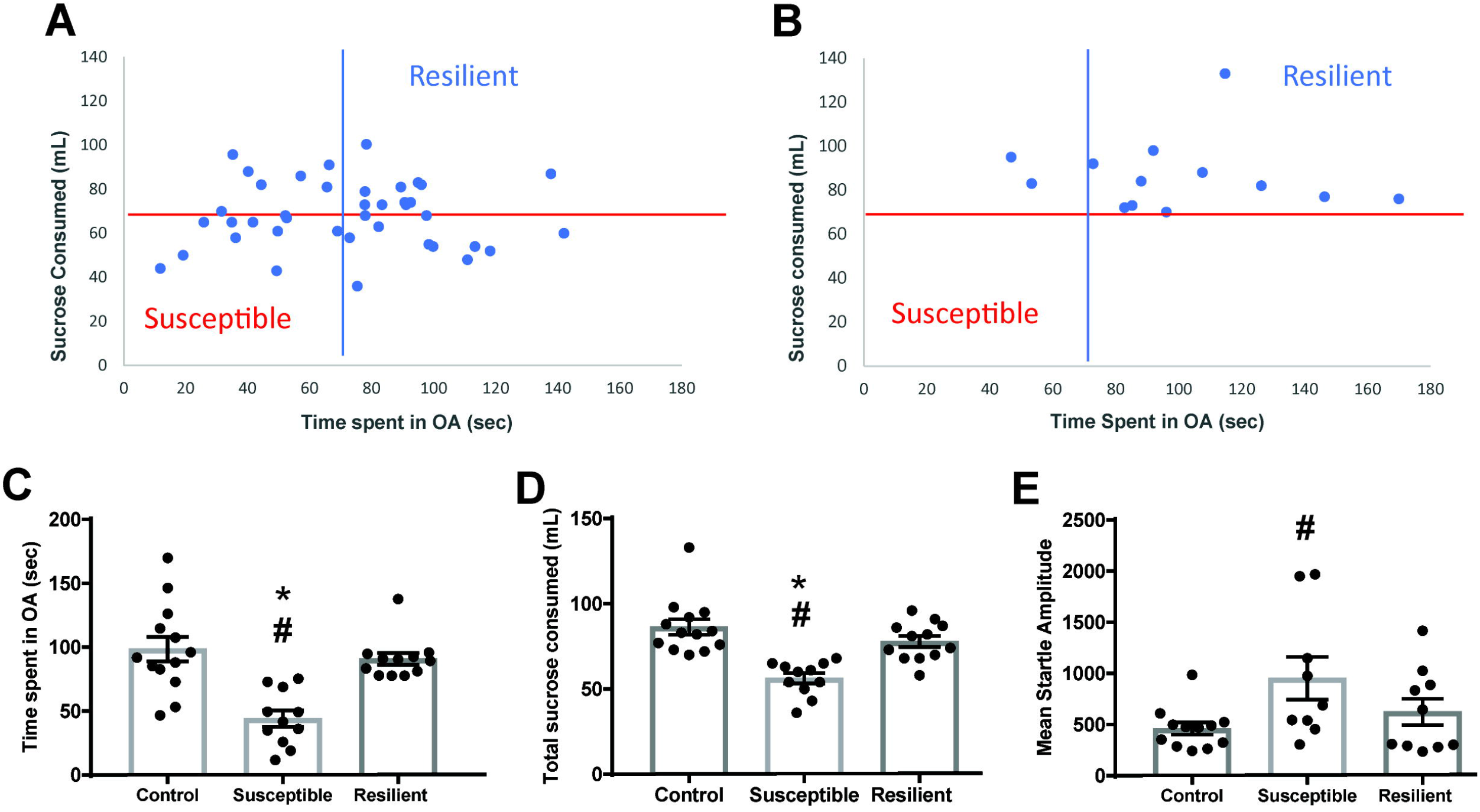
A median split analysis conducted on sucrose intake and time spent in the OA reveals stress-Susceptible and stress-Resilient phenotypes. **(A)** TMT-exposed Resilient subjects exhibited sucrose consumption and time spent in the EPM OA greater than the median for TMT-exposed rats. TMT-exposed Susceptible ats were defined as those exhibiting less than the median total sucrose and time spent in the EPM OA. **(B)** The same median split analysis conducted on control rats yielded the majority of rats exhibiting behavior similar to TMT-exposed Resilient rats and no Susceptible rats. **(C)** When rats were phenotyped based on the double-median split, this produced a Resilient group that did not differ from the Controls on time spent in the EPM OA. Susceptible rats spent less time in the OA than both Resilient and Controls. **(D)** TMT-exposed Resilient phenotype consumed similar amounts of sucrose solution as Controls; Susceptible rats consumed less sucrose than Controls and Resilient. **(E)** Mean ASR amplitude did not differ between Resilient and Controls; Susceptible rats exhibited greater startle amplitude than Controls. # = p<0.05 vs. Control; * = p<0.05 vs. Resilient.

### 3.3 Light-Dark Box

Time spent in the dark compartment of the light-dark box differed by Phenotype [F (2, 33) = 8.323, p = 0.0012, Fig. 3A]. Susceptible rats spent a greater amount of time in the dark side than Control and Resilient rats. Latency to enter the dark side did not differ between phenotypes.

**Fig. 3.**
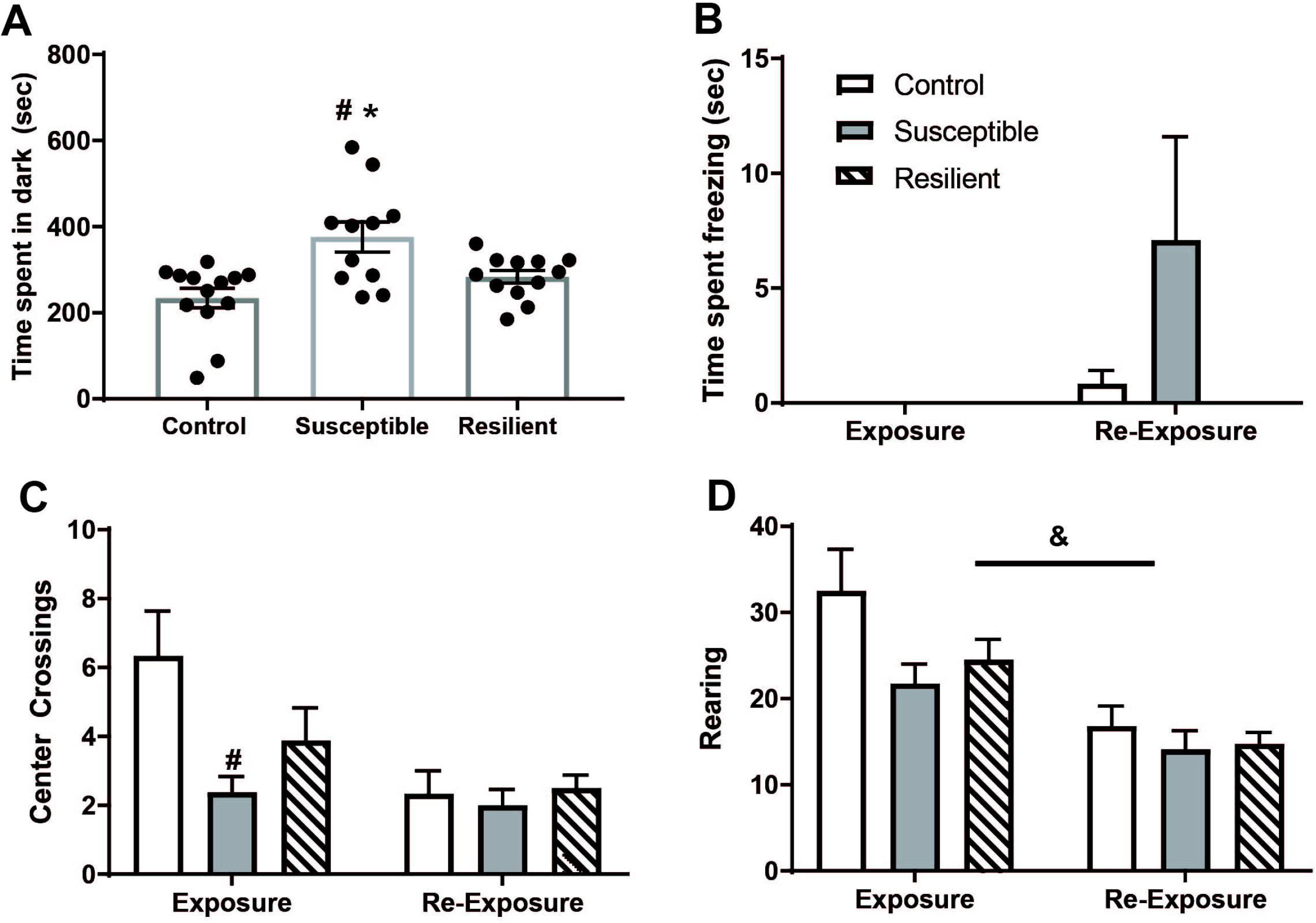
Effects of TMT-exposure on anxiety-like, conditioned fear, and active coping behaviors. **A**. The Susceptible group spentmore time in the dark side of the light-dark box relative to both Control and Resilient rats **B**. Female rats, regardless of group, did not freeze during TMT-exposure or re-exposure to the TMT chamber. **C**. Susceptible rats displayed fewer center crossings during TMT-exposure, when the TMT was directly under the center of the mesh floor, but not during the re-exposure. **D**. There was an effect of time on rearing, with all groups displaying greater rearing during the exposure than re-exposure. # = p<0.05 vs. Control. & = p<0.05 Exposure vs. Re-exposure.

### 3.4 Contextual Fear

Control, stress-Susceptible and -Resilient rats were re-exposed to the TMT context with no TMT present for a conditioned fear test. A two-way ANOVA comparing time spent freezing during exposure and re-exposure for the three phenotypes found no main effects of Time or Phenotype on freezing and no significant interaction (Fig. 3B). In fact, during TMT exposure, none of the female rats exhibited freezing, and during the contextual re-exposure, only the Control and Susceptible rats exhibited any quantifiable freezing. We also examined time spent rearing and grooming, and number of center crossings during the exposure and re-exposure, finding a significant Phenotype x Time interaction for center crossings [F (2, 19) = 4.934, p=0.0188, Fig. 3C]. Post-hoc tests found that Susceptible rats displayed fewer center crossings during the predator scent exposure only (TMT was placed below the center of the circular chamber). For time spent rearing (Fig. 3D), there was a significant effect of Time [F (1, 38) = 27.14, p<0.0001] and Phenotype [F(2, 38) = 3.367, p=0.045], but no Phenotype x Time interaction.

One-way ANOVA’s comparing the mean ASR, time spent in OA, and sucrose consumed between rats at different stages of the estrous cycle at the time of TMT exposure revealed no effect of estrous phase on these dependent measures. No relationship between estrous phase at time of testing was observed.

### 3.5 mGlu5 mRNA expression

Brain tissues from a subset of rats were used for FISH analysis; one-way ANOVAs were conducted to confirm that the same effect of phenotype was observed for this subset of rats [time spent in OA: F (2, 21) = 16.76, p<0.001; mean ASR: F (2, 17) = 7.903, p =0.0037; sucrose consumed F (2, 21) = 16.12, p<0.001]. In accordance with prior observations that the great majority of mGlu5 mRNA was expressed in vGluT1-expressing nuclei, mRNA expression in IL, PL, NRe, and BLA were contained within or immediately surrounding vGluT1-positive neuronal bodies (33).

#### 3.5.1 Prefrontal cortex

Prelimbic prefrontal cortex (PL) mRNA expression significantly differed between groups [F(2,19) = 10.51, p=0.0008; Fig. 4A], with Holm-Šídák’s post-hocs indicating both Susceptible and Resilient animals expressed greater PL mRNA than controls. Infralimbic prefrontal cortex (IL) mRNA expression did not differ between groups [F(2,18) = 0.6253, p=0.5463; not shown].

**Fig. 4.**
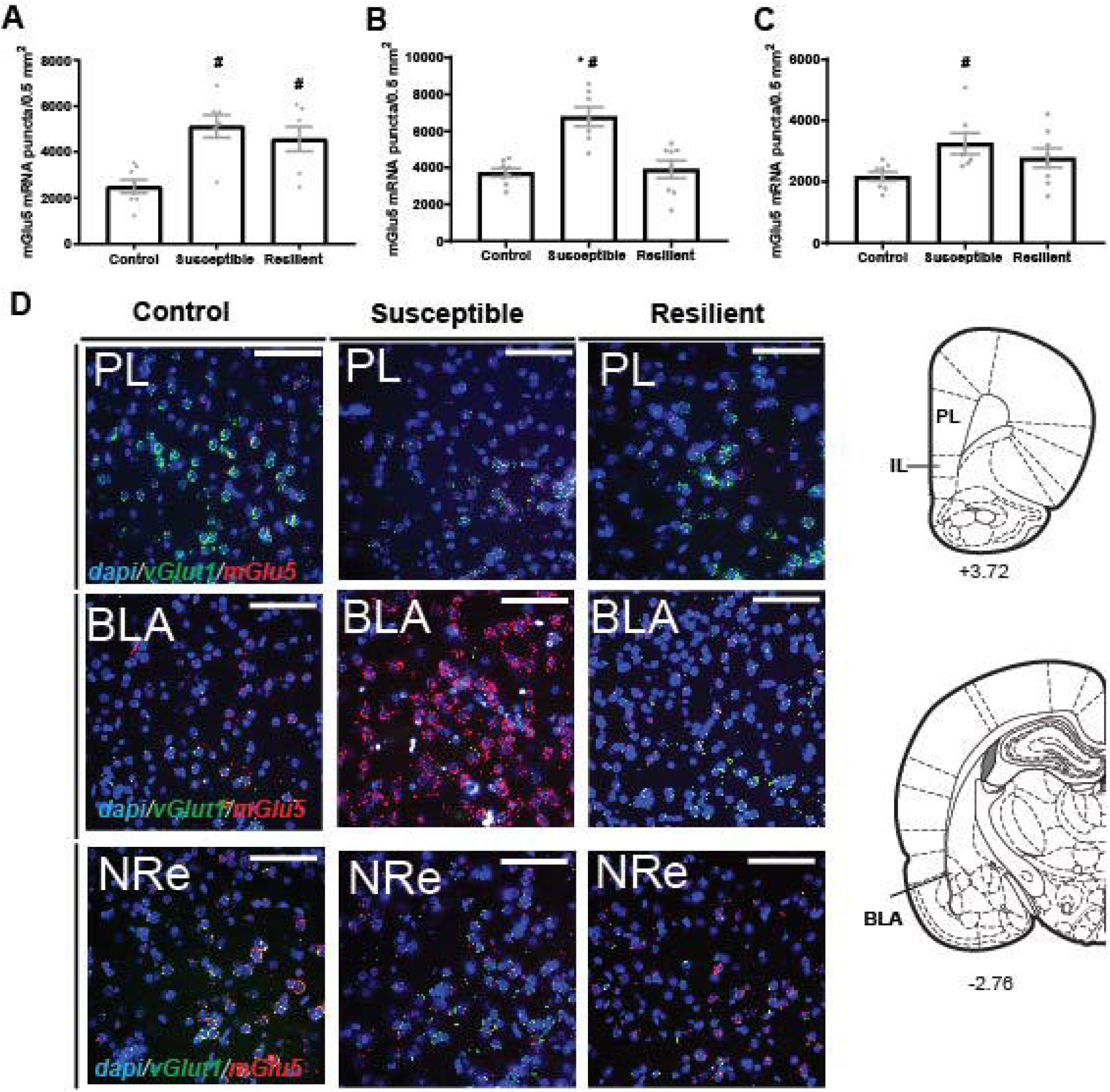
Phenotypic differences in mGlu5 expression were observed in several brain regions. **A**. mGlu5 expression in PL vGluT+ cells was greater in both Susceptible and Resilient rats relative to Controls. **B**. BLA mGlu5 expression in vGluT+ cells was greater in Susceptible rats relative to both Controls and Resilient rats. **C**. Nucleus reunion mGlu5 expression in vGluT+ cells was greater in Susceptible rats relative Controls. **D**. Representative images and sites of mGlu5 analysis. # = p<0.05 vs. Control; * = p<0.05 vs. Resilient.

**Fig. 5.**
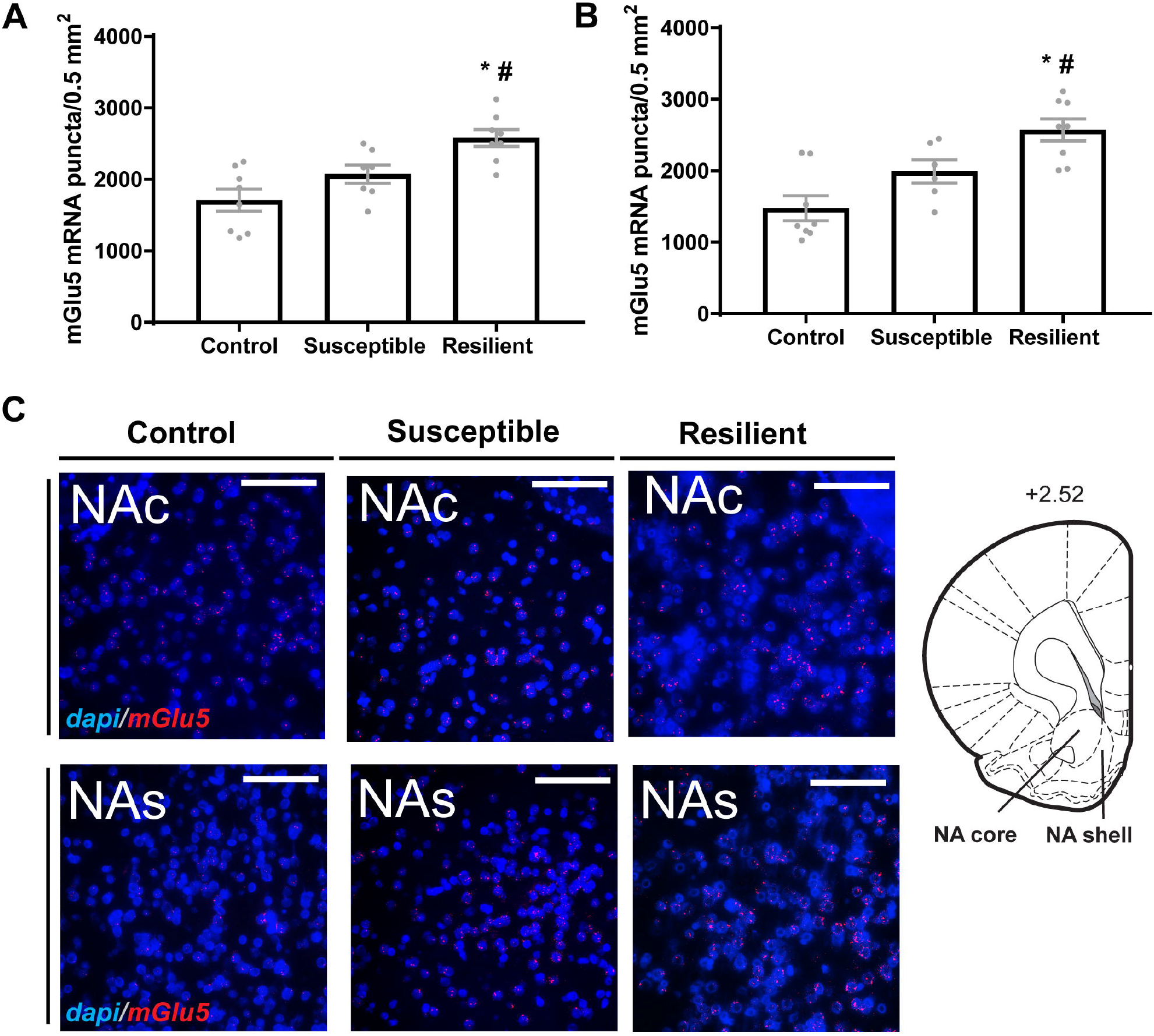
Resilient rats displayed greater mGlu5 expression in the nucleus accumbens. **A**. Nucleus accumbens core and shell **(B)** mGlu5 expression was greater in Resilient rats compared to both Control and Resilient rats. **C**. Representative images and sites of mGlu5 analysis. # = p<0.05 vs. Control; * = p<0.05 vs. Susceptible.

#### 3.7.2 Nucleus reuniens & basolateral amygdala

Stress-susceptible females exhibited elevated expression of mGlu5 mRNA in BLA when compared to controls and resilient rats [F(2,20) = 15.62, p<0.0001; Fig. 4B]. Holm-Šídák tests found that Susceptible rats displayed mGlu5 mRNA expression that was significantly greater than control (p=0.0002) and Resilient (p=0.0002) groups.

There was a significant effect of Group on NRe mGlu5 expression [F(2,20) = 3.690, p=0.0433; Fig. 4C]. Holm-Šídák’s post-hoc tests determined Susceptible rats expressed greater mRNA expression than controls (p=0.0433).

#### 3.7.3 Nucleus accumbens

We found a significant effect of group on NAcore mGlu5 mRNA expression [F(2,20) = 10.92, p=0.0006]. Holm-Šídák’s indicated that Resilient subjects expressed significantly greater NAcore mGlu5 transcripts than Susceptible or Controls. There was also a significant effect of group on NAshell mGlu5 mRNA expression [F(2,19) = 11.87, p = 0.0005]. Unlike the former, although, Holm-Šídák’s post-hoc testing indicated that Resilient females expressed greater NAshell mGlu5 mRNA expression than Controls (p=0.0003), but not Susceptible.

## 4. Discussion

The objective of this study was to determine if, like males (33,34), female rats would exhibit long-term anxiety-like behavior and fear after a single PSS (TMT) exposure. In males, the combination of reduced time spent in the open arms of the EPM and attenuated habituation of the startle response have been classically used to characterize stress Susceptible rats. Here we found that TMT-exposed female rats display greater mean startle amplitude but habituate this response akin to controls. Still, female rats display similar decrease in the time spent in the OA after TMT-exposure as do male rats in our hands. TMT also induced long-term anhedonia in females, evidenced by decreased sucrose preference and consumption. Thus, while finding distinct behavioral differences in the habituation of the ASR and contextual fear, we found that females exhibit long-term anxiety and anhedonia after TMT. Applying double median split criteria to the two variables affected by TMT (sucrose intake and time spent in the OA), yielded distinct Resilient and Susceptible populations. When considering phenotype, differences in behavior emerged, including time spent in the dark of the light-dark box and behavior during the TMT-exposure (crossing over the center area above the TMT) was reduced in rats later classified as Susceptible.

Two papers that directly compared the anxiety-like behavior of female and male Sprague-Dawley rats after a single exposure to PSS (cat urine), found that cat urine induced less anxiety-like behavior in both the EPM and ASR in females (46,47). PSS decreased time spent in OA and increased mean startle amplitude similarly in males and females; however, whether habituation of this response occurred in either sex was not discussed. Here we found that, unlike male rats, female TMT-exposed rats habituate the startle response over 30 trials. Prior clinical (48) and preclinical data (49,50) suggest that trauma-exposed females do not show analogous increase in startle responses as males. When applying cut-off behavioral criteria to segregate rats into Susceptible and Resilient phenotypes, akin to the current study, a lower incidence of Susceptible phenotype was found in female rats (15.8%) compared to males (40%) (47). While these authors concluded that female rats are more resilient to trauma than male rats, the present data indicate that the ASR model does not accurately assess female stress responses. In these studies (46,47), and in the present data set, estrous cycle phase at time of EPM/ASR testing and PSS had no effect on anxiety behaviors.

It is unclear why only a subpopulation of trauma-exposed individuals develop PTSD and why the rates are different for males and females. Some evidence suggests that circulating and de-novo estrogen and progesterone can heighten emotional memory consolidation, which in turn may contribute to higher PTSD susceptibility in women (51–57). Here we did not find any relationship between phase of the estrous cycle and the development of anxiety-like behavior. Another source of variability might be the fact that PTSD symptoms are somewhat sexually dimorphic (58–63). For example, women report affective disturbances and anhedonia, including clinical depression, at greater rates than males (64) (65). Here we found that TMT-exposed rats consume less sucrose solution and display a reduced sucrose preference compared to Controls. The decreased sucrose preference is a well-established correlate of anhedonic state in rodents that is accompanied by increased ICSS thresholds and can be reversed by antidepressants but not by neuroleptics or anxiolytics (66). Thus in our PSS model produces long-term anhedonia, similarly to what has been observed after other types of stress exposure in male rats (66–68).

Typically, in male PTSD research, freezing is a well-documented innate defensive fear mechanism (69). High amounts of freezing in re-exposure testing are thought to suggest evidence that animals exhibit re-experiencing-like symptoms. Unlike TMT-exposed Susceptible male rats, TMT-exposed female rats did not display overt freezing behavior here. In classic fear conditioning studies utilizing foot shock, males have been found to freeze more than females during fear conditioning and extinction (70) while others find the same degree of freezing between the sexes (71). Another study has reported that a subpopulation of female rats exhibits escape-like, active fear response known as “darting” during conditioned tone presentations (71). Similarly, in the forced swim test, female rats are more likely to engage in active coping strategies (72). Thus, the lack of freezing exhibited here is further evidence that freezing is not a fear response consistently exhibited by female rats. The finding here that rats later classified as Susceptible displayed fewer crossings over the center part of the apparatus where the TMT was placed indicates that they were aware of and avoided the stressor during exposure. Female Long Evans (LE) rats exhibit significant amounts of freezing during TMT exposure and during TMT-context re-re-exposure two weeks after TMT exposure (73,74). When considering that the LE strain serves as a rodent model of HPA-hyperactivity, greater paraventricular nucleus of the hypothalamus CRH-receptor expression and lower hippocampal glucocorticoid receptor expression in male LE compared to male SD rats (75) may contribute to why greater rates of freezing have been observed after TMT in female LE rats. In a direct comparison of LE to SD rats, female SD rats exhibit reduced immobility in a forced swim test relative to female LE rats. Another explanation for the previously observed freezing during TMT context re-exposure is that in this case, rats were exposed to TMT and tested during the light phase of the light cycle, where here all procedures were conducted during the dark phase. In fact, our procedures were conducted within 2 hours of the beginning of the dark phase, the time at which corticosterone levels are at their highest in unstressed Sprague-Dawley rats. Such diurnal fluctuations in corticosterone may have interacted with the TMT exposure to produce different responses in the respective studies (76).

Using the identical PSS model, we have previously assessed mGlu5 mRNA levels using the qRT-PCR approach in tissue punches, finding that Resilient male rats exhibit greater mGlu5 mRNA in the amygdala and PFC than Control and Susceptible rats. We then used FISH to determine that, relative to Controls, mGlu5 was upregulated in the PL, IL, and BLA but not the central nucleus of the amygdala. The present finding that both Susceptible and Resilient female rats exhibit elevated expression of mGlu5 in the PL is not necessarily in contradiction to our findings in males, as we never assessed Susceptible rats in a subregion-specific manner. Following a single exposure to TMT, male LE rats exhibit stress-reactive behaviors and also show upregulated mGlu5 in the PL cortex (13). Thus, independent of phenotype, the present results in TMT-exposed females are in agreement with our previous findings.

Here we found distinct differences in female BLA mGlu5 expression compared to our previous findings in males: only Susceptible female rats display greater mGlu5 expression. In male Resilient rats (but not Controls), infusion of the mGlu5 antagonist MTEP into the BLA increases freezing during re-exposure to the context (33). The nucleus reuniens is implicated as a key hub in thalamocortical integration and is involved in contextual fear in female rats exposed to TMT (74,77). We found that Susceptible rats displayed increased NRe-localized mGlu5 expression. Thus, in light of the established role of BLA mGlu5 in mediating contextual fear in males, the lack of freezing observed here in combination increased BLA mGlu5 only in Susceptible rats indicates that the neurobiology of contextual fear expression is different in female and male rats. It is possible that females exposed to TMT are engaged in active coping mechanisms (as opposed to freezing), and this behavior differentially recruits mGlu5 in the BLA and NRe.

Resilient rats displayed greater NAcore mGlu5 mRNA expression. In mGlu5 knockout mice, but not wild-type, chronic restraint stress leads to decreased sucrose preference, indicating that mGlu5 confers resilience to the long-term effects of stress (29). However, in unstressed alcohol-preferring rats, operant self-administration of sucrose is unaffected by intra-accumbens infusion of the mGlu5 antagonist MPEP (78). Thus, our work and that of others support the idea that mGlu5 is engaged by stress to regulate anhedonic processes.

## 5. Conclusions

The present work finds that female Sprague-Dawley rats display long-term anxiety-like behavior following a single exposure to TMT, and that such behavior occurs in a heterogeneous manner. However, there are distinct differences between male and female rats in this model, including the ability of females to habituate the startle response and not develop conditioned freezing to the PSS environment. Future studies will address a possible causal role for mGlu5 in the NAc in the resilience to stress-induced anhedonia in this PSS model.

## Acknowledgments

This work was funded by the University of Florida Center for OCD, Anxiety, and Related Disorders pilot grant awarded to MS.

## Declaration of Competing Interest

The authors declare that they have no conflict of interest.

## Notes

### Competing Interest Statement

The authors have declared no competing interest.

